# Embryonic origins of forebrain oligodendrocytes revisited by combinatorial genetic fate mapping

**DOI:** 10.1101/2024.01.23.576886

**Authors:** Yuqi Cai, Zhirong Zhao, Mingyue Shi, Mingfang Zheng, Ling Gong, Miao He

## Abstract

Multiple embryonic origins give rise to forebrain oligodendrocytes (OLs), yet controversies and uncertainty exist regarding their differential contributions. We established intersectional and subtractional strategies to genetically fate map OLs produced by medial ganglionic eminence/preoptic area (MGE/POA), lateral/caudal ganglionic eminences (LGE/CGE) and dorsal pallium. We found that, contrary to the canonical view, LGE/CGE-derived OLs make minimum contributions to the neocortex and corpus callosum, but dominate piriform cortex and anterior commissure. Additionally, MGE/POA-derived OLs, instead of being entirely eliminated, make small but sustained contribution to cortex with a distribution pattern distinctive from those derived from the dorsal origin. Our study provides a revised and more comprehensive view of cortical and white matter OL origins, and established valuable new tools and strategies for future OL studies.

## Introduction

Oligodendrocytes (OLs) are an important class of macroglia responsible for producing the myelin sheaths that insulate and protect neuronal axons. Forebrain OLs arise from multiple embryonic origins. Previous fate mapping study using *Nkx2.1^Cre^*[1], *Gsh2^Cre^*[2] and *Emx1^Cre^*[3] reported consecutive and competing waves of OLs derived from medial ganglionic eminence/preoptic area (MGE/POA), lateral/caudal ganglionic eminences (LGE/CGE) and dorsal pallium[2]. The first-wave of OLs generated by MGE/POA (_MP_OLs) were believed to be eliminated postnatally, while those from the second and third waves (_LC_OLs and dOLs) survive and populate the cortex and corpus callosum at comparable proportions. Several other studies provided both supporting and contradicting evidence to this model[4–11]. Moreover, *Gsh2* was recently found to be expressed in dorsal progenitors[12], casting doubt on the interpretation of lineage tracing data from *Gsh2^Cre^*.

In this study, we generated new genetic tools and combinatorial fate mapping strategies which allow direct visualization and comparison among OLs derived from different origins. We found that neocortical OLs are primarily composed of dOLs, rather than similar proportions of _LC_OLs and dOLs. In contrast, _LC_OLs and dOLs made comparable contributions to piriform cortex. We also found that although _MP_OLs only make a small contribution, they do persist in the cortex beyond adulthood with a unique spatial pattern distinct from that of the dOLs. In the two major white matter commissure tracts, dOLs are the vast majority in corpus callosum but make little contribution to anterior commissure, while _LC_OLs behaved the opposite. These findings significantly revised the classical view and provided a new and more comprehensive picture of cortical and white matter OL origins.

## Results and Discussion

To unambiguously track OLs from different embryonic origins, we first generated a knock-in driver, *Opalin^P2A-Flpo-T2A-tTA2^* (Figure 1), orthogonal to Cre drivers that label dorsal or ventral progenitors (*Progenitor^Cre^*). *Opalin* (also known as *Tmem10*) encodes oligodendrocytic myelin paranodal and inner loop protein that are specifically expressed in differentiated oligodendrocytes[13–16]. In *Opalin^P2A-Flpo-T2A-tTA2^*, *Flpo* and *tTA2* were inserted before the STOP codon and linked by self-cleavage peptide P2A and T2A (Figure 1A-C), allowing co-transcription and translation with *Opalin*. Flp-mediated recombination by this driver (hereinafter referred to as *Opalin^Flp^*for simplicity) enables highly specific, efficient and irreversible OL labeling, while the tTA2 component offers the flexibility for OL-specific labeling in tunable densities (Figure 1D-F).

**Figure 1.**
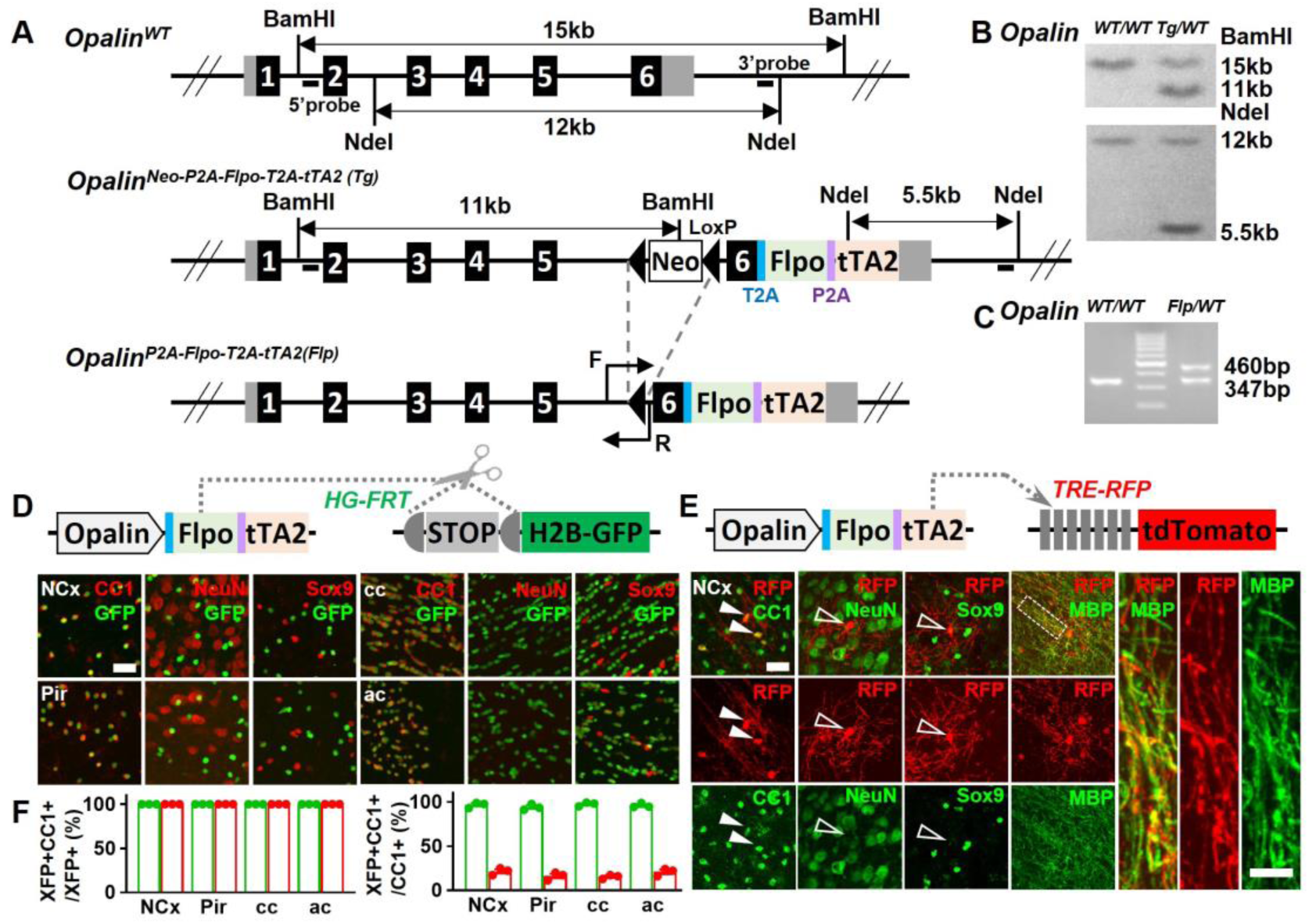
A new driver mouse for efficient and specific OL labeling. (**A**) Scheme for generating the *Opalin^P2A-Flpo-T2A-tTA2^*allele. **(B)** Southern blot confirmation of correctly targeted ES clone. **(C)** Genomic PCR to genotype F1 offspring. **(D)** OL labeling by Flp. **(E)** OL labeling by tTA2. High magnification images of the boxed region showing co-localization of RFP with MBP staining, which further demonstrated the myelination ability of labeled OLs. **(F)** Quantification of labeling specificity (left panel) and efficiency (right panel) by colacalization with OL marker CC1. Both reporting systems are highly specific, as shown by the complete co-localization of fluorescent protein (XFP) with OL marker (CC1) and lack of co-staining with neuronal marker (NeuN) or astrocyte marker (Sox9). Quantification bar-graph was not presented for NeuN and Sox9 as zero co-localizations were observed in all analyzed regions. Close to complete OL labeling was achieved by Flp-dependent H2B-GFP reporter in all analyzed regions (green dots), while sparser labeling with variable regional density was achieved by tTA2-dependent tdTomato reporter driven by TRE promoter (red dots). NCx: neocortex. Pir: piriform cortex. cc: corpus callosum. ac: anterior commissure. Scale bar: 50 μm in low magnification images, 5μm in high magnification images. Quantification: n=3. Dots represent data from individual mice.

Next, we established two types of genetic combinatorial fate mapping strategies to directly visualize OLs from different embryonic origins (Figure 2): (1) combining *Opalin^Flp^* and *Progenitor^Cre^* with intersectional reporters *Ai65* to label OLs derived from Cre+ progenitor domain by RFP (Figure 2A); (2) combining *Opalin^Flp^* and *Progenitor^Cre^* with *RC::FLTG*[17] to simultaneously label OLs derived from Cre+ progenitors by GFP and OLs derived from the complementing Cre-progenitors by RFP (Figure 2B, Supplementary Figure 1A). The first approach allowed us to track dOLs and _MP_OLs (Figure 2C, E). The second approach empowered us to observe and compare OLs generated from dorsal and ventral origins (Supplementary Figure 1B), or those from Gsh2+ and Gsh2-progenitors, in the same brain (Supplementary Figure 1C). Importantly, the subtraction power enabled us to target OLs derived from LGE/CGE progenitors that express neither *Emx1* nor *Nkx2.1* (Figure 2D and Supplementary Figure 1D). In addition, these strategies greatly facilitated the identification of OLs derived from specific origin which exist at relatively low density in certain regions.

**Figure 2.**
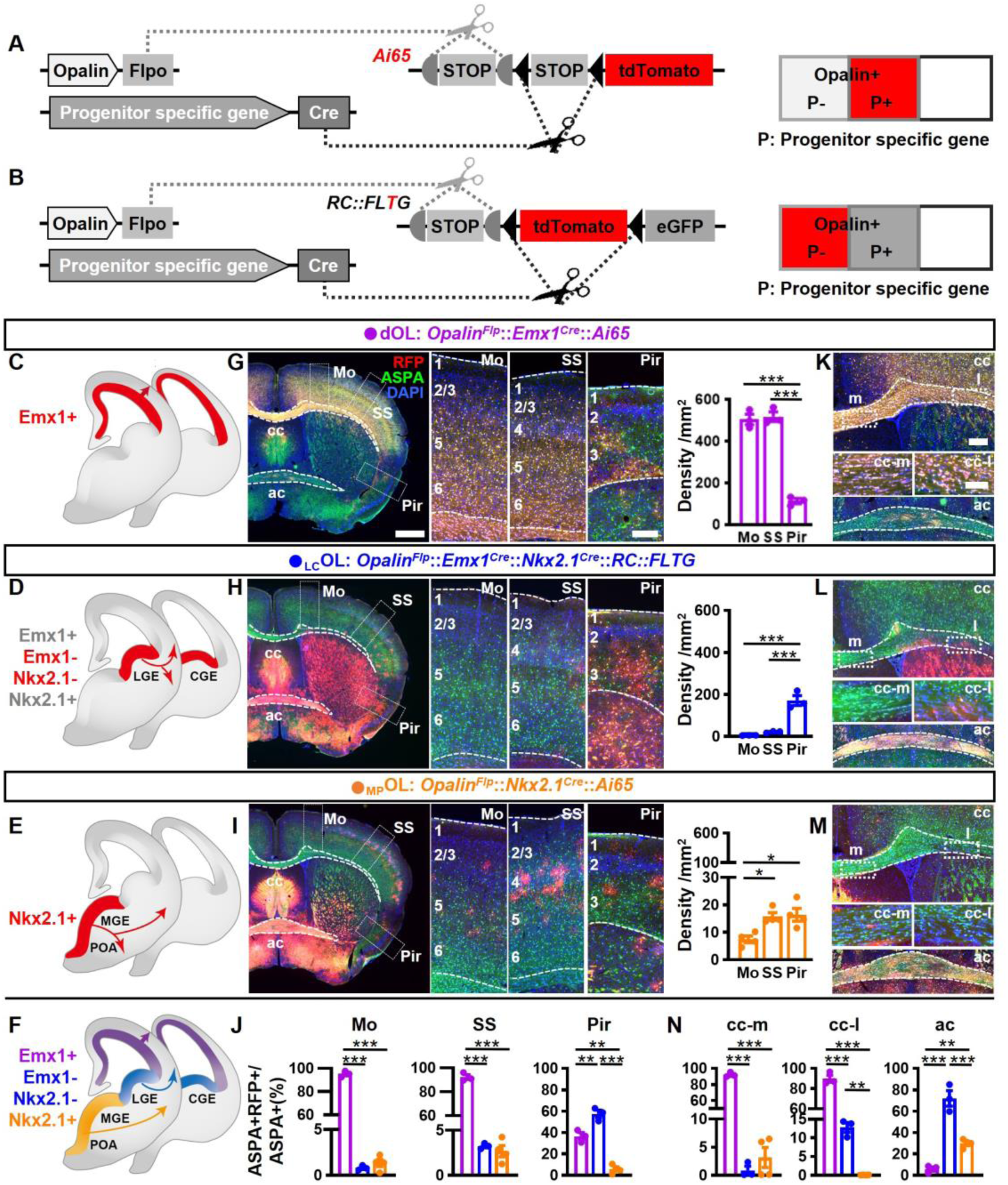
Combinatorial fate mapping of dOLs, _MP_OLs and _LC_Ols. (**A**) Strategy for intersectional labeling. Flp-AND-Cre labels OLs from Cre-expressing progenitors with RFP. **(B)** Strategy for subtractional labeling of OLs derived from non-Cre-expressing progenitors with RFP. The eGFP expressing OLs derived from Cre expressing progenitors were not used for analysis in this scenario and thereby were not highlighted by color. **(C-F)** Schematics showing intersectional labeling of dOLs in *Opalin^Flp^*::*Emx1^Cre^*::*Ai65* (C), subtractional labeling of _LC_OLs in *Opalin^Flp^*::*Emx1^Cre^*::*Nkx2.1^Cre^*::*RC::FLTG* (D), intersectional labeling of _MP_OLs in *Opalin^Flp^*::*Nkx2.1^Cre^*::*Ai65* (E) and cortical OLs derived from all three origins (F). **(G-I)** Representative images (left panels) and quantifications (right panels) of RFP+ cell density in motor cortex (Mo), somatosensory cortex (SS) and piriform cortex (Pir). **(J)** Quantification of differential contribution to ASPA+ OLs by three embryonic origins to Mo, SS and Pir. **(K-N)** Representative images (K-M) and quantifications (N) of differential contribution to ASPA+ OLs by three embryonic origins in the two major commissure white matter tracts: corpus callosum(cc) and anterior commissure (ac). _MP_OLs and _LC_OLs preferentially reside in the medial and lateral cc (cc-m and cc-l), respectively. Scale bar: 1mm in low magnification images in (G-I), 250 μm in high magnification images of the boxed area in (G-I) and low magnification images in (K-M), 100 μm in high magnification images of the boxed area (cc-m and cc-l) in (K-M). n=3 for dOLs and _LC_OLs; n=4 for _MP_OLs. Dots represent data from individual mice. Error bar: S.E.M. *P < 0.05, **P < 0.01, ***P < 0.001.

Deploying these strategies, we assessed the differential contributions of dOLs, _LC_OLs and _MP_OLs by analyzing RFP+ cells in the following mice: *Opalin^Flp^*::*Emx1^Cre^*::*Ai65* (Figure 2C), *Opalin^Flp^*::*Emx1^Cre^*::*Nkx2.1^Cre^*::*RC::FLTG* (Figure 2D) and *Opalin^Flp^*::*Nkx2.1^Cre^*::*Ai65* (Figure 2E). To better assess their contributions to the total OL population (Figure 2F), we co-stained RFP with the mature OL marker aspartoacylase (ASPA)[18] (Figure 2G-I) and quantified the ratio of colocalization (Figure 2J). Notably, all RFP+ cells are ASPA+, reassured the specificity of our label strategies. We observed two significant differences from the traditional model in the neocortex. The first major deviation is that, instead of comparable contributions by dOLs and _LC_OLs, the vast majority of neocortical OLs were dOLs but not _LC_OLs. The densities (Figure 2G-I and Supplementary Figure 2A-F) and ASPA ratios (Figure 2J) of dOLs are much higher than those of _LC_OLs. Considering the possibility of incomplete recombination in combinatorial reporters, and the relatively low Cre activity in the dorsal MGE of *Nkx2.1^Cre^*[1], the genuine contribution of _LC_OLs to the neocortex could be even lesser than our current observation. Therefore, the large quantity of neocortical OLs labeled by *Gsh2^Cre^*in previous study[2] or by GFP in *Opalin^Flp^*::*Gsh2^Cre^*::*RC::FLTG* (Supplementary Figure 1C) most likely were predominantly dOLs generated by Gsh2+ dorsal progenitors[12], rather than bona fide _LC_OLs.

The second major deviation is that cortical _MP_OLs are not completely depleted postnatally. Instead, they make a small but continued contribution with a unique spatial distribution pattern (Figure 2I-J and Supplementary Figure 2D-G). _MP_OLs display a clear rostrocaudal density decline (Supplementary Figure 2E-F), a higher density in somatosensory cortex (SS) than motor cortex (Mo) (Figure 2I), and a laminar preference towards layer 4 (L4) in SS (Supplementary Figure 2G). In contrast, the distribution of dOLs and _LC_OLs do not vary significantly across the rostrocaudal axis (Supplementary Figure 2E-F) or between Mo and SS (Figure 2G-H), but exhibits increased density towards deeper layers (Supplementary Figure 2G). Importantly, we have observed cortical _MP_OLs in mice as old as one year (Supplementary Figure 2H), well beyond the age analyzed in previous reports[2, 7, 8], suggesting a persisted contribution.

We then turned our attention to the lateral three-layer archicortex, piriform cortex (Pir). Different from the neocortex, Pir contains higher proportions (Figure 2J) of _LC_OLs than dOLs. _MP_OLs make the lowest contribution (Figure 2J) at a density similar to SS and higher than Mo (Figure 2I).

These combinatorial models also grant us the opportunity to revisit the differential contributions of dOLs, _LC_OLs and _MP_OLs to the two commissural white matter tracts, corpus callosum (cc) and anterior commissure (ac), which contain high density of OLs (Figure 2K-M). We found that, similar to the neocortex, cc is mainly populated by dOLs and supplemented by very low proportions of _LC_OLs and _MP_OLs (Figure 2K-N). Interestingly, _LC_OLs and _MP_OLs seem to show preferential distribution in the lateral and medial regions of cc (cc-l and cc-m), respectively (Figure 2L-N). Different from cc, ac is mainly populated by _LC_OLs and _MP_OLs and supplemented by very low proportion of dOLs (Figure 2K-N).

To substantiate the above results, we further breed *Opalin^Flp^*::*Emx1^Cre^*::*Nkx2.1^Cre^*::*Ai65* to label dOLs together with _MP_OLs by RFP and co-stained them with ASPA (Supplementary Figure 3). RFP-ASPA+ cells were difficult to find in Mo, SS and cc, but were more easily observed in Pir and ac, consistent with the respective low and high _LC_OL contributions in these regions.

In summary, our findings significantly revised the canonical model of forebrain OL origins (Figure 3A), and provided a new and more comprehensive view (Figure 3B). We demonstrated that neocortical OLs are mainly derived from dorsal origin with small but lasting contribution from the ventral origin (Figure 2, Supplementary Figure 1B and 2). Our data showed that LGE/CGE makes little contribution to neocortex and cc, but makes major contribution to piriform cortex and ac (Figure 2 and Supplementary Figure 3). This finding is supported by another report in which in utero electroporation failed to label LGE-derived cortical OLs in both embryonic and early postnatal brains, and an exclusion strategy revealed very low percentage of LGE/CGE-derived cortical OLs in neonatal brains[19]. The lack of adult labeling in our study together with the lack of developmental labeling in the other study suggest that the lack of _LC_OL in neocortex is less likely caused by competitive postnatal elimination, but more likely due to limited production and/or allocation. We further discovered that MGE/POA makes a small but persistent contribution to the neocortex with a distinct distribution pattern featured by a rostral-high to caudal-low gradient and a preference towards L4 in SS (Supplementary Figure 2). Whether their enduring existence and highly biased localization has functional implications awaits future exploration. In addition, we found that the cc showed a similar OL composition as the neocortex, but the Pir and the ac each exhibited distinct OL compositions in term of their embryonic origins. _LC_OLs are the major contributor to both regions, while dOLs and _MP_OLs mainly contribute to Pir and ac, respectively (Figure 2).

**Figure 3.**
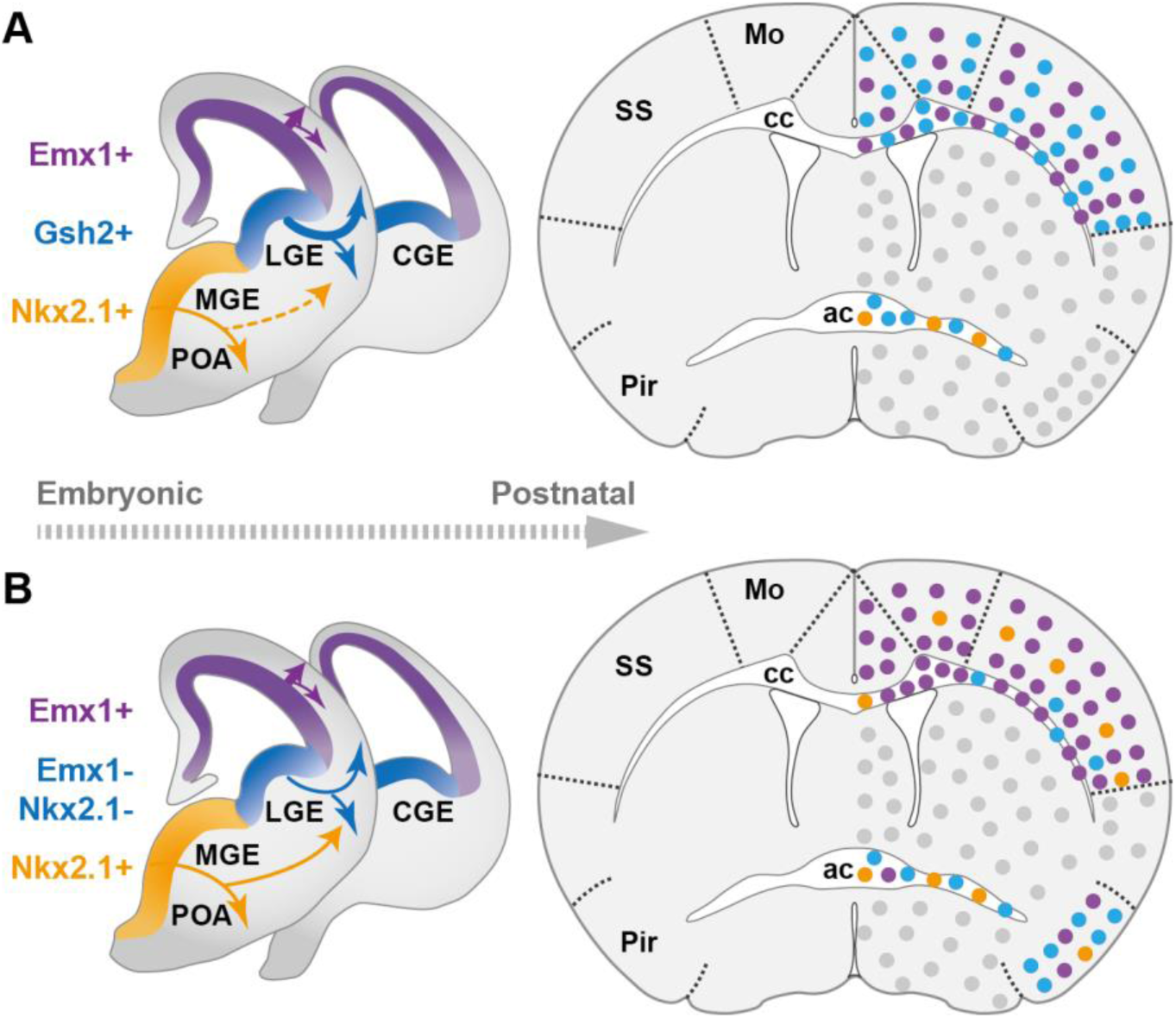
The classical and revised model of forebrain OL origins. (**A**) In the classical model[2], OLs derived from MGE/POA (orange) were largely eliminated postnatally (thin dashed line), while those from LGE/CGE (blue) and dorsal origin (purple) survive at similar proportions (thick solid line). Therefore, neocortex (NCx) and corpus callosum (cc) contain comparable density of _LC_OLs (blue dots) and dOLs (purple dots) and are devoid of _MP_OLs (orange dots). **(B)** In the new model, NCx and cc mainly contains dOLs with very low contribution from the ventral origins. _LC_OLs mainly contribute to piriform cortex (Pir) and anterior commissure (ac). _MP_OLs makes a small but sustained contribution to NCx, with a strong laminar preference towards layer 4 in somatosensory cortex (SS). In addition, dOLs and _MP_OLs also make substantial contributions to Pir and ac, respectively. Grey dots indicate OLs in unanalyzed regions.

In addition to the new framework of forebrain OL origins (Figure 3), we also generated a new driver (Figure 1) and established multiple combinatorial genetic models (Figure 2) for efficient tracking and direct visualization of OLs from different embryonic origins without interference from other cells types sharing the same progenitor domains such as oligodendrocyte precursors, astrocytes, and neurons (Figure 1-2). These tools set up a firm foundation and will provide reliable experimental access for future inquiries on the development and function of diverse OLs in healthy and disease brains[20], especially to uncover the relationship between their developmental origins and the functional and molecular heterogeneity.

## Material and Methods

### Mice

All mouse studies were carried out in strict accordance with the guidelines of the Institutional Animal Care and Use Committee of Fudan University. All husbandry and experimental procedures were reviewed and approved by the same committee. All applicable institutional and/or national guidelines for the care and use of animals were followed. The following transgenic mouse lines were used in this study: *Nkx2.1^Cre^* (Jax 008661)[1],*Gsh2^Cre^*(Jax 025806)[2], *Emx1^Cre^* (Jax 005628)[3], *Ai65* (Jax 021875)[21], *RC::FLTG* (Jax 026932)[17]. The tTA2-dependent tdTomato reporter (*TRE-RFP*) was derived from *Ai62* (Jax 022731) [21], by removing LoxP-STOP-LoxP with *E2a-Cre* (Jax 003724). The Flp dependent H2B-GFP reporter (*HG-FRT*) was derived from *HG-dual* (Jax 028581) via removal of loxP flanking STOP cassette by *CMV-Cre*[22]. The *Opalin^P2A-Flpo-T2A-tTA2^* allele was generated by targeted insertion of the T2A-Flpo-P2A-tTA2 sequence immediately before the STOP codon of the endogenous *Opalin* gene using homologous recombination. Gene targeting vector was generated using PCR-based cloning approach as described before[22]. More specifically, a 4.7 kb 5’ homology arm, a loxP flanking Neo positive selection cassette, a T2A-Flpo-P2A-tTA2 cassette and a 2.7 kb 3’ homology arm were cloned into a building vector containing the DTA negative selection cassette to generate the targeting vector. Targeting vector was linearized and transfected into a C57/black6 ES cell line. ES clones that survived through negative and positive selections were first screened by genomic PCR, then confirmed by Southern blotting using appropriate DIG-dUTP labelled probes. One positive ES cell clone was used for blastocyst injection to obtain male chimera mice carrying the modified allele following standard procedures. Chimera males were bred with C57BL/6J females to confirm germline transmission by genomic PCR. The Neo selection cassette was self-excised during spermatogenesis of F0 chimeras. Heterozygous F1 siblings were bred with one another to establish the colony. Targeting vector construction, ES cell transfections and screening, blastocyst injections and chimera breeding were performed by Cyagen.

### Genomic PCR

Genomic DNA was prepared from mouse tails. Tissue was lysed by incubation in tail lysis buffer (Viagen, 102-T) with 0.1 mg/ml proteinase K (Diamond, A100706) overnight at 55°C followed by 45 min at 90°C in an air bath to inactivate proteinase K. The lysate was cleared by centrifugation at maximum speed (21,130 G) for 15 min in a table-top centrifuge. Supernatant containing genomic DNA was used as the PCR template for amplifying DNA products. The following primers were used:

*Opalin-F*: 5’-GGCCTATGTTTGATTTCCAGCACTG –3’

*Opalin-R*: 5’-AGCACTTATGACTGCTGAGCCGTTC –3’

### Immunohistochemistry and microscopy

Mice were anaesthetized by intraperitoneal injection of 1.5% sodium pentobarbital (0.09 mg/g body weight) and then intracardially perfused with saline followed by 4% paraformaldehyde in 0.1 M PB. Following post fixation at 4°C for 24 hours, brain samples were sectioned at 30 μm using a vibratome (Leica VT1000S), or transferred into 30% sucrose in 0.1M PB for cryoprotection, OCT embedded, and sectioned using a cryostat (Leica CM1950). For CC1 immunostaining, antigen retrieval was performed prior to blocking by boiling for 3 min in 10 mM citrate buffer (pH6.0). Sections were blocked in PBS containing 0.05% Triton and 5% normal donkey serum and then incubated with the following primary antibodies in the blocking solution at 4 ℃ overnight: RFP (goat polyclonal antibody, 1:2000, SICGEN AB0081-200; rabbit polyclonal antibody, 1:2000, Rockland 600-401-379), GFP (chicken polyclonal antibody, 1:1000, Aves Labs, GFP-1020), MBP (rat polyclonal antibody, 1:500, AbD Serotec, MCA409S), CC1 (rabbit polyclonal antibody, 1:500, Oasis Biofarm, OB-PRB070, mouse polyclonal antibody, 1:300, Millipore, OP80), ASPA (rat polyclonal antibody, 1:200, Oasis Biofarm, OB-PRT005), Sox9(rabbit polyclonal antibody, 1:2000, Chemicon, AB5535), NeuN (mouse monoclonal antibody, 1:500, Millipore, MAB377). Sections were then incubated with appropriate Alexa fluor dye-conjugated IgG secondary antibodies (1:500, Thermofisher Scientific) or CF dye-conjugated IgG secondary antibodies (1:250, Sigma) in blocking solution and mounted in Aqua-mount (Southern Biotech, 0100-01). Sections were counterstained with DAPI. Sections were imaged with confocal microscopy (Olympus FV3000), fluorescence microscopy (Nikon Eclipse Ni; Olympus VS120; Olympus VS200) and fluorescent stereoscope (Nikon SMZ25). All quantifications were performed in 2-month-old adult mice from coronal sections between Bregma +1.94 and –2.80 mm. Anatomical regions were identified according to the *Paxinos “The Mouse Brain” Atlas* and the *Allen Reference Atlas*, and their areas were measured in ImageJ for density calculations, whenever applicable. For cortical regions, every fourth section within the range of selection was analyzed. For whiter matter tracts, three consecutive sections at Bregma 0.14 were analyzed. At least three brains were analyzed for each genotype. To quantify density and co-localization, cells were identified and counted in Adobe Photoshop or Image J in conjugation with QuPath.

### Statistical analysis

GraphPad Prism version 8.0.1 was used for statistical calculations. No statistical methods were used to predetermine sample sizes, but our sample sizes are similar to those reported in previous publications. Data collection and analysis were performed blind to the conditions of the experiments whenever possible. No animals or data points were excluded from the analysis. Normalcy was assessed using Shapiro-Wilk test. Equal variances were assessed using F test or Bartlett’s test. Statistical significance was tested using two-tailed unpaired t-test, Welch’s t-test, one-way ANOVA and two-way ANOVA followed by Tukey’s or Bonferroni post hoc test, wherever appropriate. Data are presented as mean ± standard error of the mean (S.E.M.). P < 0.05 was considered significant. Significance is marked as *P < 0.05; **P < 0.01 and ***P < 0.001.

## Acknowledgements

We thank Dr. Yilin Tai for helpful discussion and Dr. Min Jiang from core facility of IOBS, Fudan University, for technical support on imaging experiments. This study was supported by funds from the National Science and Technology Innovation 2030 Major Projects of China (STI2030-Major Projects-2022ZD0206500), National Natural Science Foundation of China (32171087, 32200918, 32371145).

## Author contributions

Conceptualization: MH

Methodology: YC, MH

Investigation: YC, ZZ, MS

Visualization: YC, MH, LG

Supervision: MH, LG

Writing—original draft: MH, YC, LG

Writing—Review and editing: MH, YC, ZZ, MZ

## Supplementary information

**Supplementary Figure 1.**
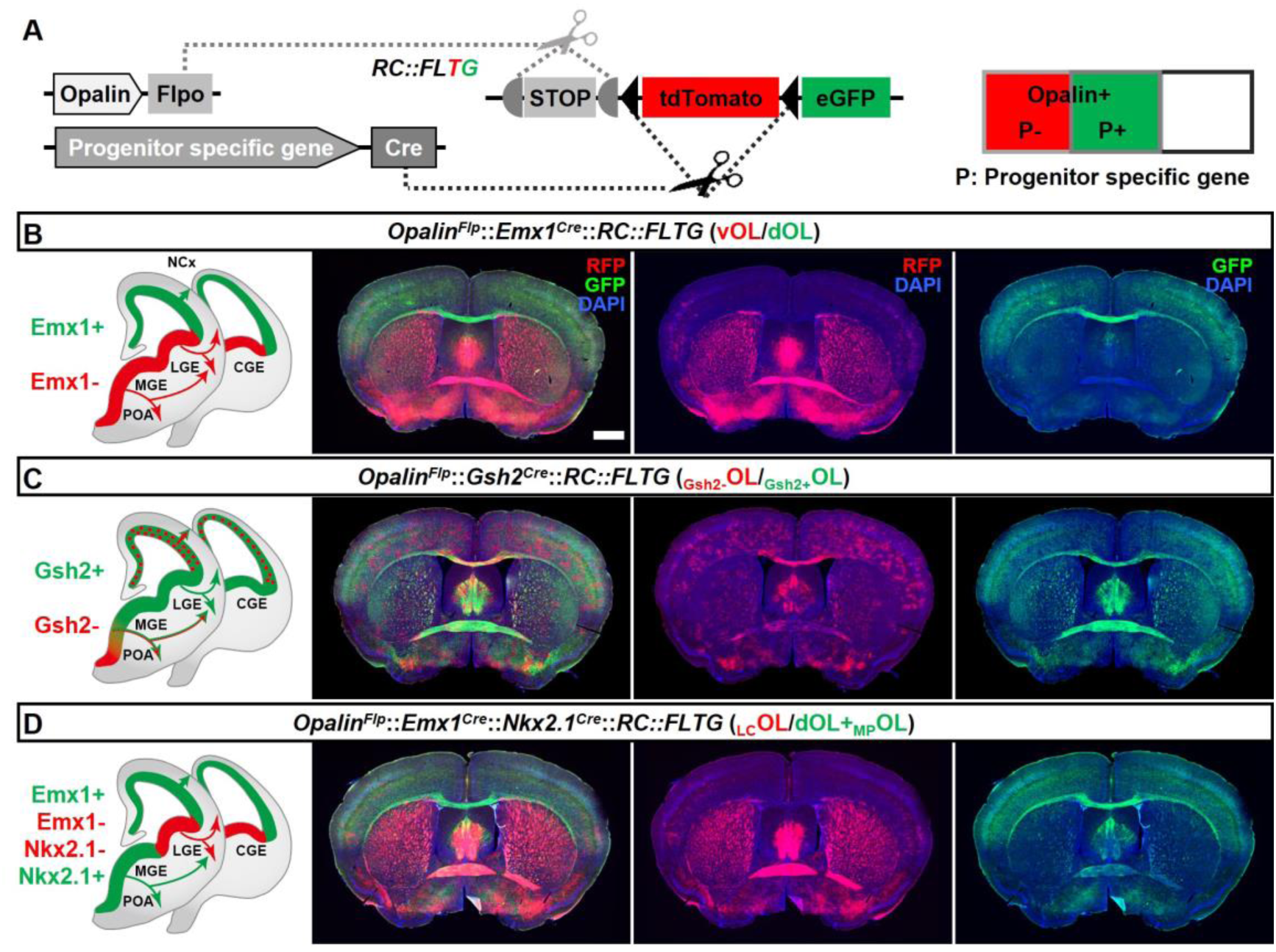
Simultaneous differential labeling of OLs derived from complementary embryonic origins. (**A**) Strategy for simultaneous labeling of OLs derived from complementary origins. Flp-NOT-Cre labels OLs from non-Cre-expressing progenitors with RFP, while Flp-AND-Cre labels OLs from Cre-expressing progenitors with eGFP. **(B)** Coronal sections showing GFP+ OLs from dorsal origin (dOLs) and RFP+ OLs from ventral origin (vOLs) in *Opalin^Flp^*::*Emx1^Cre^*::*RC::FLTG.* **(C)** Coronal sections showing GFP+ OLs derived from Gsh2+ progenitors (_Gsh2+_OL) and RFP+ OLs derived from Gsh2-progenitors (_Gsh2-_OL) in *Opalin^Flp^*::*Gsh2^Cre^*::*RC::FLTG.* **(D)** Coronal sections showing GFP+ OLs from dorsal and MGE/POA origin (dOLs+_MP_OLs) and RFP+ OLs from LGE/CGE origin (_LC_OLs) in *Opalin^Flp^*:: *Emx1^Cre^:: Nkx2.1^Cre^*::*RC::FLTG.* Scale bar: 1mm.

**Supplementary Figure 2.**
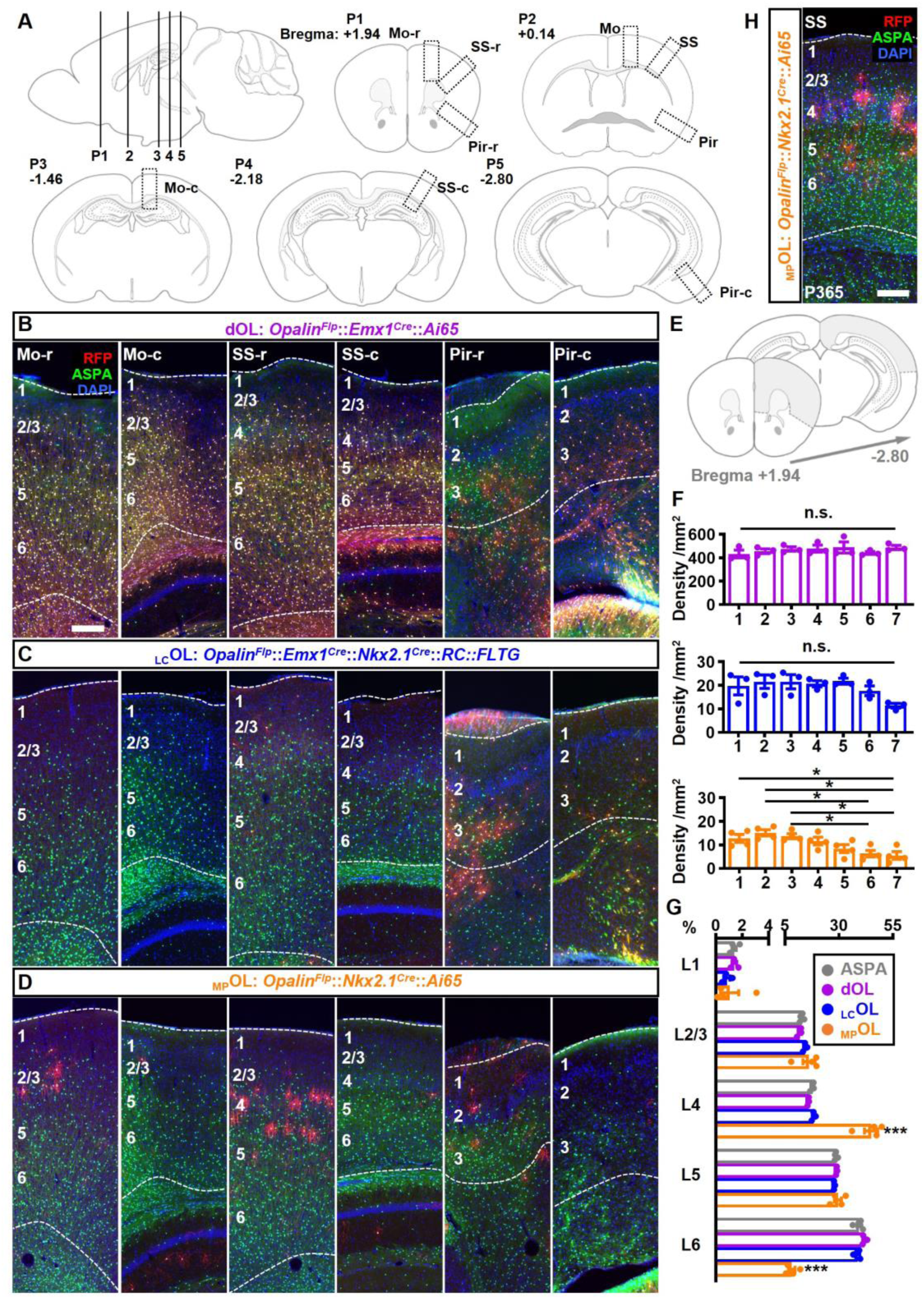
The distribution pattern of cortical dOLs, _MP_OLs and _LC_OLs. (**A**) Schematics of cortical regions chosen for quantifications and for showing representative images (boxed regions). Every fourth coronal section between Bregma +1.94 and –2.80 mm was analyzed. Position (P) 1 to 5 correspond to sections from which representative images were taken from. P2 corresponds to the sections shown in Figure 2G-I. **(B-D)** For each cortical region, two representative images at the rostral (r) and caudal(c) ends were presented for each combination. **(E-F)** Quantification of rostrocaudal distribution of neocortical (grey shaded region in E) dOLs, _LC_OLs and _MP_OLs. Slices were grouped into seven bins numbered rostrocaudally. Densities of dOLs and _LC_OLs showed no significant change across bins, while _MP_OLs exhibited lower density in more caudal regions. **(G)** Distributions of dOLs, _LC_OLs and _MP_OLs across 6 layers in SS. Similar to the total OL distribution quantified based on ASPA staining, more dOLs and _LC_OLs reside in deeper layers. In contrast, _MP_OLs are highly enriched in L4 at the cost of L6 with significant deviation from the total OLs. **(H)** Representative image of SS from 1 year old *Opalin^Flp^*::*Nkx2.1^Cre^*::*Ai65* mouse. Scale bar: 200 μm. n=3 for dOLs and _LC_OLs; n=4 for _MP_OLs and ASPA. Dots represent data from individual mice. Error bar: S.E.M. *P < 0.05, ***P < 0.001.

**Supplementary Figure 3.**
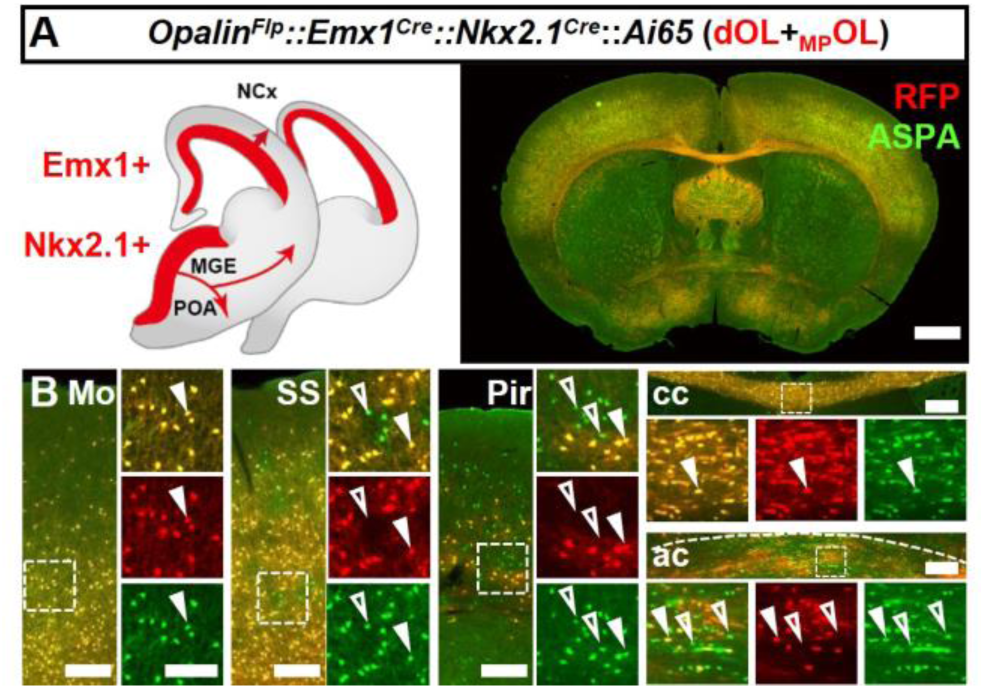
Intersectional labeling of OLs derived from both dorsal origin and MGE/POA. (**A**) Coronal sections showing both dOLs and _MP_OLs labeled by RFP in *Opalin^Flp^*::*Emx1^Cre^*::*Nkx2.1^Cre^*::*Ai65.* **(B)** Higher magnification images showing ASPA+ RFP+ dOLs/_MP_OLs (closed arrow heads) and ASPA+RFP-putative _LC_OLs (open arrow heads).The latter cells were difficult to find in neocortical regions such as motor cortex (Mo) and somatosensory cortex (SS), and corpus callosum (cc), but were frequently encountered in piriform cortex (Pir) and anterior commissure (ac). Scale bar: 1mm in low magnification images, 200μm in Mo, SS, cc and ac, 10 μm in high magnification images of boxed area.

